# Electric field-induced pore constriction in the human Kv2.1 channel

**DOI:** 10.1101/2024.12.18.629208

**Authors:** Venkata Shiva Mandala, Roderick MacKinnon

**Author notes:** Correspondence to: Roderick MacKinnon. **Email:**.

## Abstract

Gating in voltage-dependent ion channels is regulated by the transmembrane voltage. This form of regulation is enabled by voltage sensing domains (VSDs) that respond to transmembrane voltage differences by changing their conformation and exerting force on the pore to open or close it. Here we use cryogenic electron microscopy to study the neuronal K_v_2.1 channel in lipid vesicles with and without a voltage difference across the membrane. Hyperpolarizing voltage differences displace the positively charged S4 helix in the voltage sensor by one helical turn (∼5 Å). When this displacement occurs, the S4 helix changes its contact with the pore at two different interfaces. When these changes are observed in fewer than four voltage sensors the pore remains open, but when they are observed in all four voltage sensors the pore constricts. The constriction occurs because the S4 helix, as it displaces inward, squeezes the right-handed helical bundle of pore lining S6 helices. A similar conformational change occurs upon hyperpolarization of the EAG1 channel. Therefore, while K_v_2.1 and EAG1 are from distinct architectural classes of voltage-dependent ion channels, called domain-swapped and nondomain-swapped, the manner in which the voltage sensors gate their pores is very similar.

**Significance Statement:** Our ability to transmit signals across long distances rapidly – for example an instruction from the brain to the muscles in our fingers – depends on electrical impulses that travel along nerve cells. These electrical signals are mediated by membrane proteins called voltage-dependent ion channels. These channels have voltage sensors, which are domains that sense the voltage difference across the cell membrane and switch the channel on or off accordingly. Scientists discovered two architectural classes of voltage-dependent ion channels distinguished by the different ways the voltage sensors attach to the pore. This study shows that the two architectures are not very different after all because they both solve the problem of regulation of the pore by voltage sensors in the same way.

## Introduction

Voltage-dependent ion channels open or close their pores in response to changes in the membrane potential. The flow of ions through these channels in turn changes the transmembrane voltage. This feedback mechanism underlies cellular electricity, including the generation of action potentials in neurons (1, 2) and the initiation of contraction in skeletal muscle cells (3).

The function of these ion channels – whether they conduct K^+^, Na^+^, Ca^2+^ or other cations – is captured in two structural domains (4–7). One is the pore domain, comprised of two helices S5 and S6 that line the pore, and a selectivity filter that selects among different ions. The second is the voltage sensor domain (VSD) consisting of four transmembrane helices, S1 through S4. S4 contains repeats of the amino acid triplet (RXX)_n_, where the arginine (R) can be replaced by another positive charged or hydrophilic amino acid, and X denotes a hydrophobic residue. It is the displacement of this S4 helix under an applied electric field (created by a voltage difference across the membrane) (8–10), which is coupled to the pore domain in some manner, that ultimately determines whether the pore of the channel is open or closed. But how the movement of S4 is coupled to the pore domain remains the biggest outstanding question in this area, since most structures of such channels have been captured in the absence of a voltage difference (i.e. 0 mV) (7, 11). We refer to these as depolarized conformations. Determining hyperpolarized (i.e., negative inside voltage) conformations of these channels is not possible when the channels are solubilized in micelles or nanodiscs.

We recently captured hyperpolarized structures of two voltage-dependent potassium (K_v_) channels, EAG1 and KCNQ1, using polarized (negative-inside) liposomes (12, 13). These channels differ from each other in their architecture: EAG1 (K_v_10.1) is a nondomain-swapped channel, which means that the VSD of one subunit in the tetrameric channel packs against the pore domain of the same subunit (14). In this channel, which lacks an S4-S5 linker helix due to its nondomain-swapped configuration, the S4 helix itself clamps the pore shut in its hyperpolarized conformation (12). Meanwhile, KCNQ1 (K_v_7.1) is a domain-swapped channel where the VSD of one subunit contacts the pore domain of an adjacent subunit (15). In KCNQ1, which has an unusual S4-S5 linker helix compared to other domain-swapped K_v_ channels and requires the signaling lipid PIP_2_ for function, the S4 forms an extended loop in its hyperpolarized structure that occludes PIP_2_ binding rather than directly regulating the pore (13). In the present study, we ask how does the VSD to pore coupling operate in a domain-swapped channel that is not regulated by PIP_2_?

To answer this question, we conduct a similar cryogenic electron microscopy (cryo-EM) analysis using a domain-swapped K_v_ channel – the neuronal ‘delayed rectifier’ K_v_2.1 – where the pore is regulated directly by the voltage sensor. We discover that S4 changes its conformation near the intracellular membrane surface and contacts the pore through two interfaces (much akin to what we observed in the nondomain-swapped EAG1), accompanied by only a small displacement of the S4-S5 linker. The movement of S4 directly constricts the pore at its narrowest region formed by the conserved PXP motif in S6. Our findings thus suggest a unifying mechanism for voltage-dependent gating that applies to both domain-swapped and nondomain-swapped channels.

## Results

### Polarization of K_v_2.1-containing liposomes

To clarify notation, we refer to lipid vesicles in which we generate a negative-inside voltage (relative to outside) as polarized vesicles (or polarized sample or dataset) and lipid vesicles without a membrane voltage difference as unpolarized vesicles (or unpolarized sample or dataset). We refer to channel conformations driven by polarized vesicles as hyperpolarized channels and channel conformations observed in the absence of polarization as depolarized channels. We make this distinction for the following reason. When vesicles are subject to a polarizing environment (see below), it appears that only a fraction of them, presumably those that are not leaky to ions and can maintain their ion gradient, become polarized. Thus, we observe a distribution of hyperpolarized and depolarized channels in polarized samples. On the other hand, unpolarized vesicles contain only depolarized channels.

The full-length human Kv2.1 channel was purified and reconstituted into liposomes composed of 90:5:5 1-palmitoyl-2-oleoyl-sn-glycero-3-phosphocholine (POPC) to 1-palmitoyl-2-oleoyl-sn-glycero-3-phosphoglycerol (POPG) to cholesterol [wt/wt/wt] with 300 mM KCl. As we described previously for EAG1 and KCNQ1 (12, 13), valinomycin was added to the vesicles and the extravesicular solution was exchanged rapidly to 300 mM NaCl using a buffer-exchange column (**Figure 1A**). The valinomycin (or K_v_2.1) mediated K^+^-efflux generates a membrane potential with an upper limit of about −145 mV, such that the inside of the vesicles is negative with respect to the outside. These polarized vesicles were immediately applied to a holey grid and frozen for cryogenic electron microscopy (cryo-EM) analysis. As noted above, we are likely to have vesicles with a range of membrane potentials (between 0 mV and -145 mV) in this preparation, but we do not know what this distribution looks like. The unpolarized sample in this case contains symmetric KCl and should be at 0 mV.

**Figure 1.**
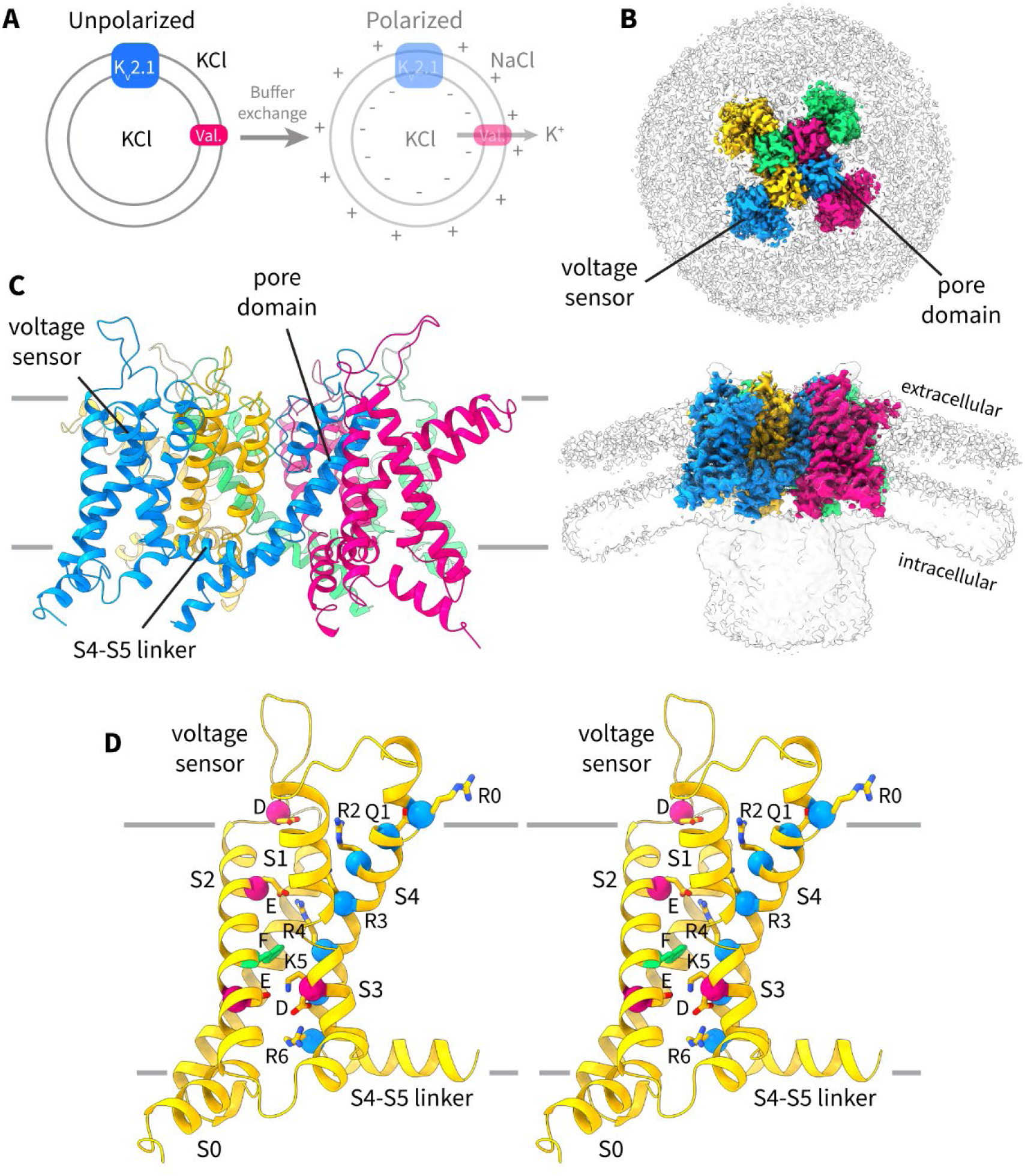
Structure of K_v_2.1 in unpolarized proteoliposomes. (**A**) Schematic of the protocol used to obtain polarized vesicles for cryogenic electron microscopy (cryo-EM) analysis. K_v_2.1-containing liposomes are prepared with symmetrical KCl and valinomycin (val.) is added to mediate K^+^-flux. The external KCl is exchanged for NaCl using a buffer-exchange column. Potassium efflux through valinomycin generates a potential difference across the membrane such that the inside of the vesicle is negative with respect to the outside. (**B**) Top-down and side views of the cryo-EM density map of the up structure of the K_v_2.1 channel from the unpolarized dataset. Each channel subunit is shown in a different color, and the lipid bilayer and the disordered cytoplasmic domain are visible at a lower contour (*grey*). (**C**) Structure of K_v_2.1 (depolarized-highK) in the unpolarized dataset (cartoon representation). Each subunit is colored differently. (**D**) Stereoview of the K_v_2.1 voltage sensor (cartoon representation) in the up conformation. The six positive charges in S4 (Cα marked by blue spheres), three negative charges in S2 and S3 (E233, S243, and D266), and the hydrophobic Phe in S2 (F240, green sticks) are shown in stick representation.

### The depolarized structure of K_v_2.1

The structure of K_v_2.1 in unpolarized vesicles (**Figure 1B, C**) and in nanodiscs has been reported previously (13, 16) and is very similar to that of other K_v_ channels such as K_v_1.2 and Shaker (17–19). In the unpolarized dataset (3.0 Å overall resolution, **Supplementary Figure S1**), the transmembrane structure is well defined while the cytoplasmic domain is only partially ordered (**Figure 1B**). The structure (**Figure 1C**) shows an open pore and voltage sensors in the ‘up’ conformation, thus being consistent with a depolarized conformation (hereby this structure is termed ‘depolarized-highK’ for reasons that will become apparent). Further 3D classification of this dataset yielded no additional classes, indicating that when we do not polarize vesicles, the channel structure is homogeneous. In other words, the voltage sensors do not move (much) of their own accord at 0 mV.

In the depolarized voltage sensor conformation (**Figure 1D**) the fifth positive-charged residue in S4, K5 (K309), resides in the gating charge transfer center comprising F240 and E243 from S2 and D266 from S3. R6 (R312) is located below the gating charge transfer center, and Q1 (Q297), R2 (R300), R3 (R303) and R4 (R306) are located on the extracellular side of the gating charge transfer center. R4 occupies the negative charged pocket on the extracellular side and interacts with E233 from S2. Notably, unlike in Shaker (or K_v_1.2) (17–19), the K_v_2.1 paddle (comprising the top parts of S3 and S4) does not pack tightly against S1 and S2, but rather bends radially out towards the lipid membrane and leaves a sizable (and presumably solvent-filled) vestibule at the extracellular side of the voltage sensor (**Figure 1C, D**). This structural difference might shape the local electric field around the voltage sensors differently in the two channels and could contribute to differences in gating charge measurements (20–22).

### A non-canonical selectivity filter conformation due to low external [K^+^]

As we have observed previously for EAG1 and KCNQ1, the polarized (**Figure 2A**) dataset was notably more heterogeneous. Through 3D classification, we identified one homogenous set of particles, corresponding to about 30% of all ‘good’ particles, that yielded a 3.3 Å reconstruction (**Figure 2B, C**). This structure (**Figure 2C**, **Supplementary Figure S2**) shows an open pore and homogeneously ‘up’ voltage sensors similar to the depolarized conformation in the unpolarized dataset (i.e. the depolarized-highK conformation) with one notable difference: the selectivity filter adopts a non-canonical, partially dilated conformation (**Figure 2C-E**). We can think of at least two reasons for this unusual conformation: 1) the low external [K^+^] (∼1 mM) in this polarized vesicle preparation (**Figure 2A**), or 2) the applied electric field. The voltage sensors of this set of particles appear to be all ‘up’ (when classifying without symmetry), indicating that these channels likely lie in vesicles that have lost (at least partially) their ion gradient, and hence their voltage difference. Thus, we ascribe the selectivity filter dilation to the low external [K^+^], noting that even if the internal solution of all the vesicles is exchanged with the outside, the external [K^+^] will still be low (i.e., the volume of the vesicles is small compared to the extravesicular volume).

**Figure 2.**
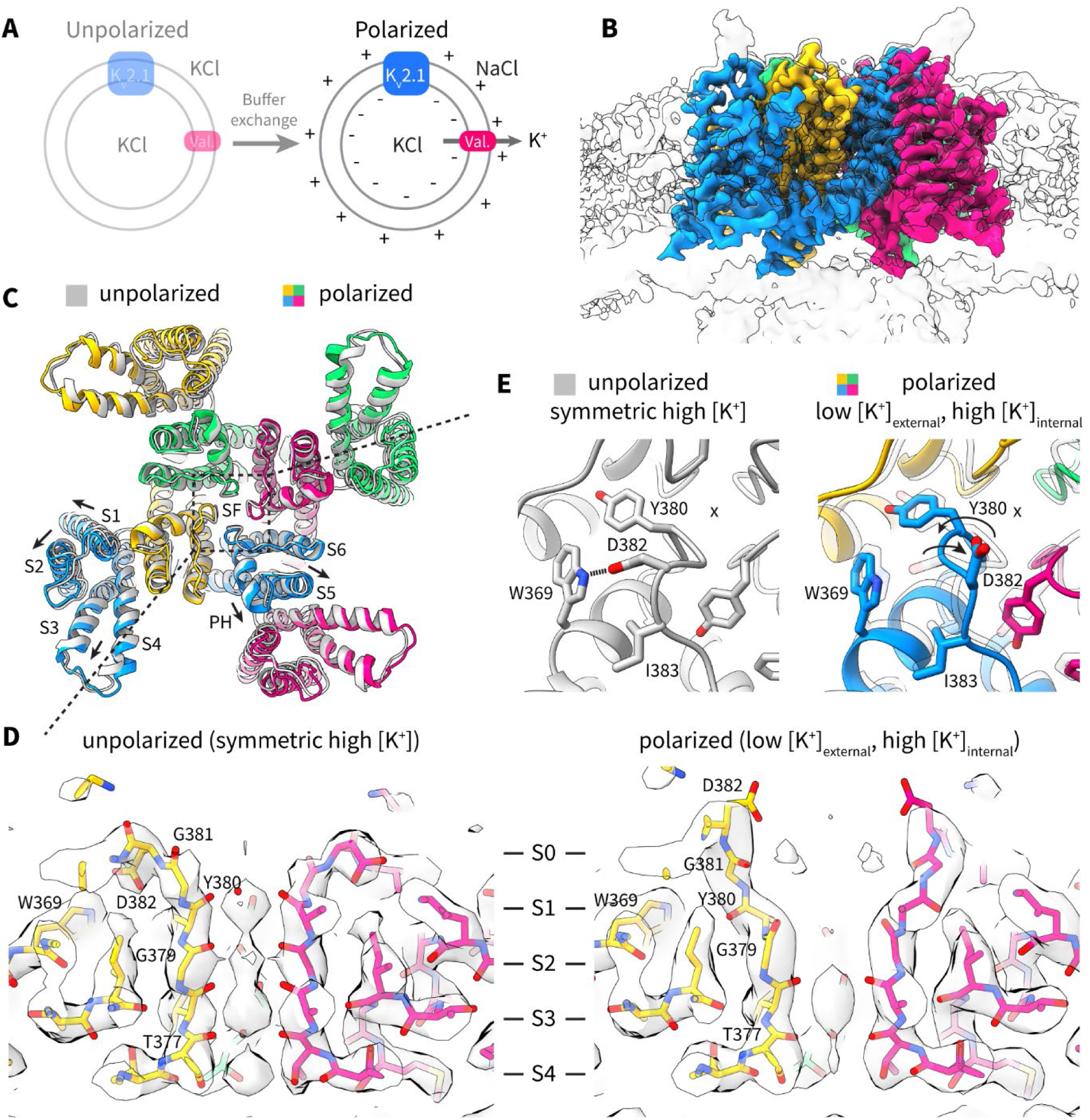
Depolarized structure of K_v_2.1 from polarized proteoliposomes. (**A**) Schematic of the vesicle-polarization protocol. (**B**) Cryo-EM density map of the depolarized structure of K_v_2.1 from the polarized dataset. Each channel subunit is shown in a different color, and the lipid bilayer is visible at a lower contour (*grey*). (**C**) Overlay of structures (cartoon representation) of depolarized K_v_2.1 in the polarized dataset (*colored*, depolarized-lowK) and in the unpolarized dataset (*grey*, depolarized-highK). (**D**) Side views of the selectivity filter of K_v_2.1 (stick representation) and the cryo-EM density (grey surface) in the unpolarized dataset (*left*) and in the polarized dataset (*right*). The positions of the four main ion binding sites (S1 through S4) and the exit site S0 are indicated. (**E**) Zoomed-in view of the selectivity filter region that undergoes a conformational change (x marks the ion permeation axis). In the unpolarized dataset structure (*left*, grey), W369 in the pore helix makes a hydrogen bond with D382 in the selectivity filter. This interaction is lost in the polarized dataset structure (*right*, colored), and G381 and Y381 rotate outward.

This non-canonical conformation is characterized by an outward rotation of the backbone carbonyls of Y380 and G381 in the selectivity filter (**Figure 2D, E**), which is accompanied by loss of the hydrogen bond between D382 near the apex of the filter and W369 in the pore helix (**Figure 2E**). Also apparent is a reduction of density for potassium ions in the selectivity filter (**Figure 2D**). Other residues in the surrounding region do not undergo significant conformational changes (**Figure 2E**). Functional implications (or the lack thereof) of this selectivity filter conformation will be discussed in a following section. The channel also shows a slight global expansion with the non-canonical selectivity filter (**Figure 2C**), indicating that the structure of the entire channel is coupled to these changes in the filter, and prompting us to use this structure (hereby termed ‘depolarized-lowK’) for comparison to the hyperpolarized structures discussed below.

### Intermediate conformation of the channel: 2 ‘down’ voltage sensors with an open pore

The remaining ∼70% particles in the dataset were heterogeneous (**Figure 3A-C**), but most appeared to have at least one voltage sensor (when classifying without symmetry) in a shifted position when compared to the depolarized conformation (**Figure 3D**). We first focus on one class (no applied symmetry) where 2 out of the 4 voltage sensors are in a ‘down’ position but the pore has remained open (**Figure 3E**), resolved to an overall resolution of 4.3 Å (**Figure 3B**, **Supplementary Figure S3**).

**Figure 3.**
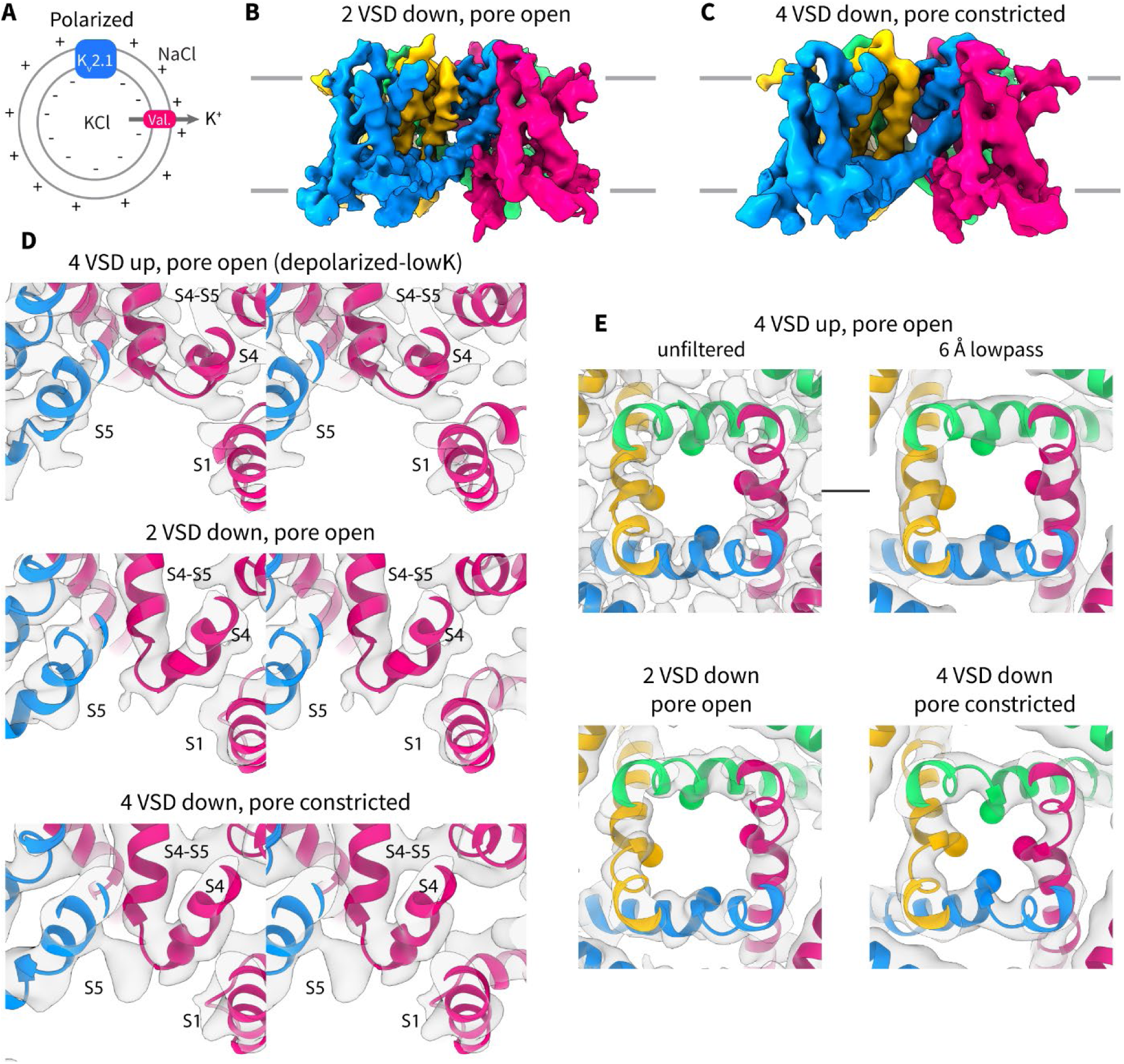
Hyperpolarized structures of K_v_2.1 from polarized proteoliposomes. (**A**) Schematic of polarized vesicles. (**B, C**) Cryo-EM density map of the (**B**) intermediate conformation with two voltage sensors down, 2 voltage sensors up, and an open pore, and of the (**C**) pore constricted conformation with four voltage sensors down. Each subunit is colored differently in the two maps. (**D**) Stereoviews of the connection between the S4-S5 linker and S4 in the depolarized (*top*), intermediate (*middle*), and constricted (*bottom*) conformations. The channel is shown in cartoon representation and the Cα of R312 is shown as a magenta sphere for reference. (**E**) Top-down view of the pore (S6) in the three conformations: depolarized model and unfiltered map (*top left*), same depolarized model with a 6Å-filtered map (*top right*), intermediate model and unfiltered map (*bottom left*), and the constricted model and unfiltered map (*bottom right*). The Cβ of P410, the second proline in the P-X-P motif, is shown as a sphere and each subunit is colored differently.

In the ‘down’ voltage sensors, the connection between the S4-S5 linker and S4 is displaced and S4 forms an extended interfacial segment (comprising A311, R312 and H313) that is directly contiguous to S5 from the adjacent subunit (**Figure 3D, Figure 4A, B**). S2 and S1 stay largely in place. The fourth positive-charged residue, R4, now occupies the gating charge transfer center. The one-helical turn (∼5 Å) downward movement of S4 is accompanied by partial loss of helical density at the top of S3 and S4 and repositioning of the S3-S4 loop. This movement is accompanied by an α-helix to 3_10_-helix transition that has been described in other channels (17, 23, 24). The 3_10_-helix region stays in place (**Supplementary Figure S4**) in the membrane while the residues that form this segment change as S4 slides down (**Figure 4A, B**). Despite the movement of two voltage sensors out of four, the pore is still open (**Figure 3E**) and the S4-S5 linker stays in place (**Figure 4B**).

**Figure 4.**
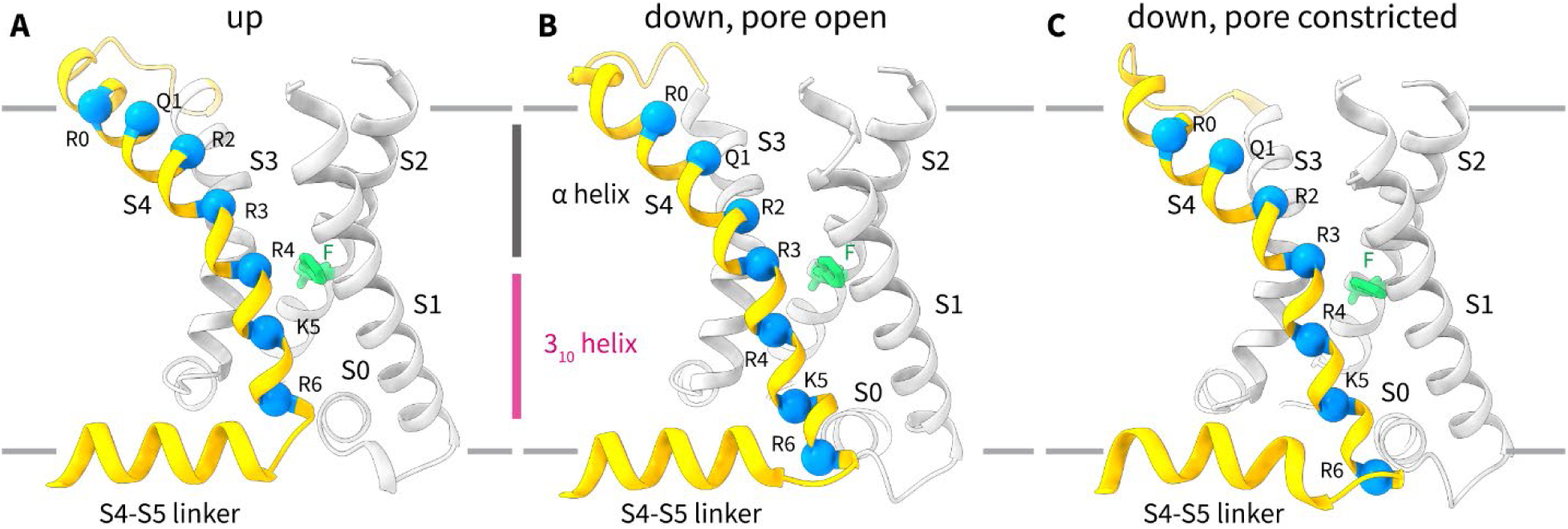
Voltage sensor conformational changes in K_v_2.1. (**A-C**) Side view of one K_v_2.1 voltage sensor in the (**A**) up conformation with an open pore, (**B**) down conformation with an open pore, and (**C**) down conformation with a constricted pore. The position of the 3_10_-helical (*magenta*) and α-helical (*dark grey*) segments in the up position and in the down position with an open pore is shown. All structures are shown in cartoon representation, with the Cα positions of the six basic residues in S4 shown as blue spheres and the gating charge transfer center residue F240 shown in green stick representation.

The selectivity filter in this class (and in other ones not discussed here) adopts the same non-canonical conformation as in the depolarized-lowK structure described above. This conformation, and other ones with one ‘down’ voltage sensor that closely resemble this structure, likely represent intermediate states of channel closing, where not a sufficient number of the voltage sensors have moved to close the pore. A discussion of these states in the context of the full gating mechanism of the channel will follow later.

### Hyperpolarized conformation of the channel: 4 ‘down’ voltage sensors and a constricted pore

Classification without alignment of a smaller subset of the ∼70% heterogeneous particles in the polarized dataset revealed lower resolution classes with constricted pores. We detail one of these classes (C4 symmetric) resolved to 5.8 Å (**Figure 3C**, **Supplementary Figure S5**) with all 4 voltage sensors in the down position (**Figure 3D**) and a constricted pore (**Figure 3E**).

The ‘down’ position of the voltage sensors is quite like that described for the intermediate class above (**Figure 3D**), at least at the lower resolution resolved for this class. S4 clearly slides down by at least 1 turn and is thus modeled as a one-helical turn movement (**Figure 4C**). Also apparent in this class is a slight downward deflection of the N-terminal end of the S4-S5 linker (**Figure 4C**) compared to the other classes (**Figure 4A, B**). We note that classification of this particle set without symmetry yielded classes where 1-2 voltage sensors appear further displaced than the others. But the lower resolution of these classes precluded unambiguous modeling of two helical turn movements. We nevertheless point out the presence of this asymmetry to suggest that the voltage sensor can move more than just one-helical turn in K_v_2.1.

While the voltage sensor movements are also visible in the intermediate class, more striking is the constriction of the pore in this class (**Figure 3E**, **Figure 5**). The conformational change is modeled as a rotation of the bottom half of S6 (S6_lower_) relative to the upper half of S6 (S6_upper_) around the conserved hinge region (**Figure 5A, D**). The hinge is formed by the P-X-P motif (X = hydrophobic residue, here isoleucine) that is conserved in most domain-swapped K_v_ channels. This rotation is in the clockwise direction when viewed from the extracellular side of the channel and results in a narrowing of the pore diameter at the bundle crossing from ∼13 Å to ∼9.5 Å (**Figure 5E**) at the narrowest rigid carbon, the Cβ of P410. If one includes hydrogen atoms, the diameter is ∼8 Å – near the diameter of a hydrated potassium ion – making it conceivable that this state represents a closed pore. Higher resolution structures will be needed to delineate the precise conformational changes that occur during pore closure.

**Figure 5.**
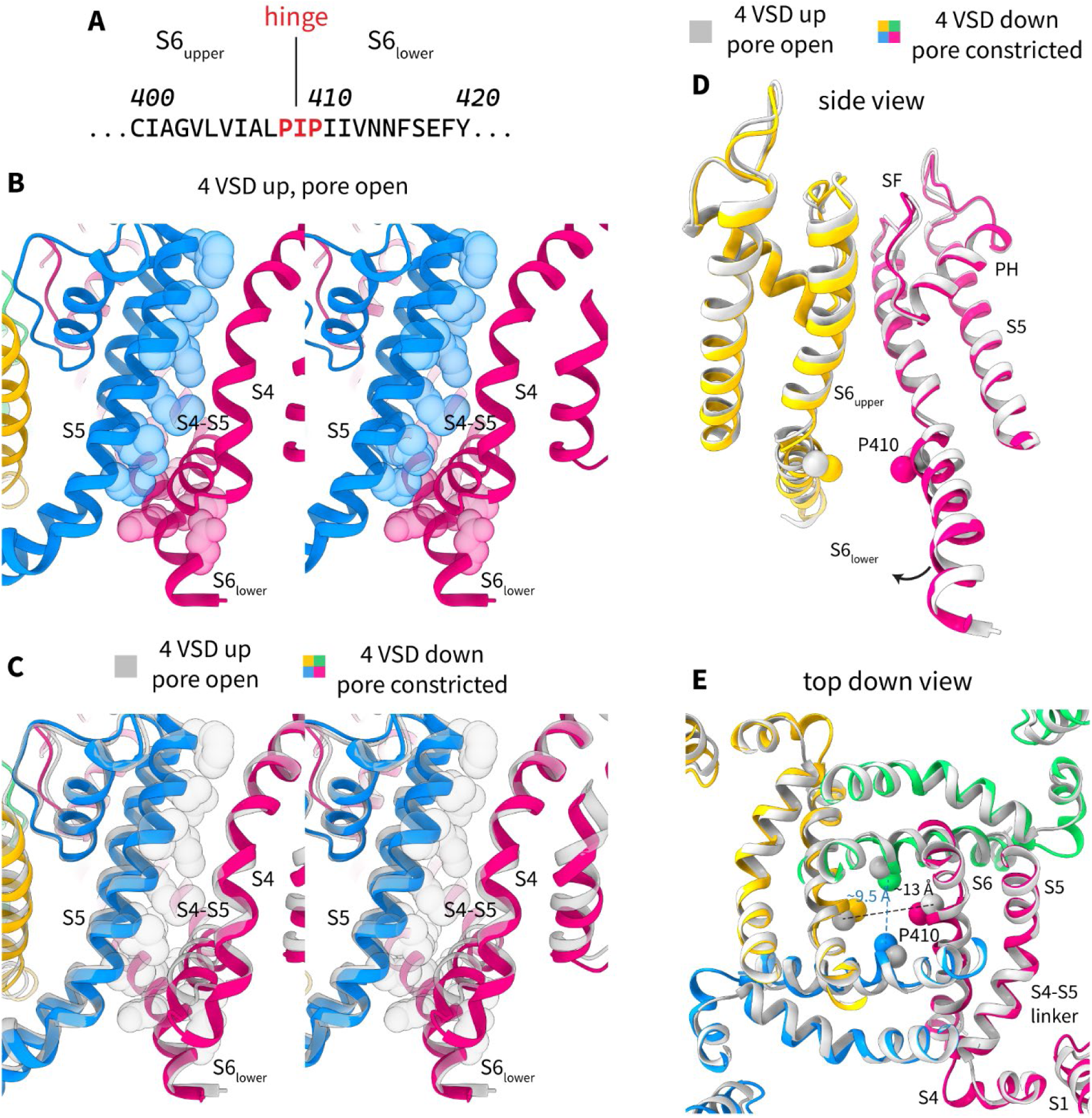
Electromechanical coupling and pore constriction in K_v_2.1. (**A**) Primary sequence of an S6 segment in K_v_2.1 showing the P-X-P hinge motif and the upper and lower parts of S6. (**B, C**) Stereoviews showing the contact between S4 in one subunit, and the S6_lower_ of the same subunit or the S5 of an adjacent subunit, in the (**B**) depolarized (*colored*) conformation and in the (**C**) depolarized (*grey*) and constricted (*colored*) conformations. The structures are shown in cartoon representation and contacting residues in the depolarized conformation are shown as spheres. (**D, E**) Side view (**D**) and top-down view (**E**) of the pore in the open conformation (*grey*) and in the constricted conformation (*colored*). The conformations are shown in cartoon representation and the Cβ of P410 is shown as spheres. In the side view, only two opposite subunits are shown for clarity.

### Coupling of voltage sensor movements to pore closure

How do the voltage sensor movements lead to pore closure? Using the well-defined depolarized-lowK structure with all four voltage sensors up and an open pore, we can examine the contacts between the S4/S4-S5 linker and the rest of the protein (**Figure 5B**). Residues on the S5 of the adjacent subunit and S6_lower_ of the same subunit that are proximal to S4 or the S4-S5 linker are shown as spheres. There are two principal interfaces: 1) the S4 of one subunit contacts the S5 of an adjacent subunit both directly and through the S4-S5 linker, and 2) the S4 of one subunit contacts S6_lower_ of the same subunit through the S4-S5 linker. Now consider what happens when the S4 moves down as membrane voltage is applied (**Figure 5C**). The S4, as it moves down (also see **Movie S1** and **Movie S2**), would apply a force directly on the pore through S5 and on S6_lower_, causing the pore constriction described above. There are thus at least two pathways for electromechanical coupling in this type of channel. Readers who are familiar with our recent work on the non-domain swapped channel EAG1 will note striking similarities between the two channels – this will be discussed further below.

## Discussion

### Non-canonical selectivity filter: related to channel inactivation or merely the resting state?

In the unpolarized sample with symmetric high [K^+^], the selectivity filter adopts a canonical, conductive conformation with ion density visible for the four binding sites. In the polarized sample with low external [K^+^] and high internal [K^+^] (in the fraction with hyperpolarized voltage sensors), the selectivity filter adopts the non-canonical conformation described earlier (**Figure 2C-E**), where the first two ion binding sites are abrogated while the bottom two are intact. Other non-canonical selectivity filter conformations have been observed before in other potassium channels. It has been shown that symmetric low [K^+^] can either induce dilation (Shaker, K_v_1.2, K_v_1.3) (18, 19, 25, 26) or constriction of the selectivity filter (hERG, KcsA) (27–29). But the case presented here constitutes an unusual selectivity filter conformation under physiological conditions of low external [K^+^] and high internal [K^+^]. In addition, the dilated conformation observed here in K_v_2.1 differs from those reported in Shaker, K_v_1.2 and K_v_1.3, constituting a much smaller conformational change (only three residues are involved) that has the same outcome of abolishing the first two ion binding sites.

Given that low external [K^+^] is known to promote C-type inactivation in some potassium channels (30, 31), it is tempting to speculate that the selectivity filter conformation we observe under these conditions is related to inactivation in K_v_2.1. Indeed, we did not observe an alternate conformation of the selectivity filter in EAG1 and KCNQ1 (12, 13) when subject to the same asymmetric ionic conditions – and these two channels do not inactivate. We cannot exclude this possibility, but the low external [K^+^] condition observed here occurs naturally in cells at the resting membrane potential. Thus, the selectivity filter of K_v_2.1 would likely adopt this non-canonical conformation under resting conditions, where the low external [K^+^] destabilizes the canonical conformation. The selectivity filters of different potassium channels might simply require varying concentrations of potassium ions to adopt their canonical conformation.

In fact, even if the selectivity filter conformation is strictly dictated by the external [K^+^], it is unclear why the outward passage of potassium ions through a canonical filter (as would occur during a depolarization event) would then eventually lead to the non-canonical conformation (i.e. when inactivation would occur). Perhaps the structure of the selectivity filter is coupled to the structure of the voltage sensor or the pore (as we indeed observe here), leading to different structures under changing membrane potentials. We thus remain agnostic about selectivity filter-mediated inactivation in K_v_2.1 (16), but merely highlight the presence of an easily accessible, external [K^+^]-dependent, non-canonical conformation of the selectivity filter in this channel.

### Electromechanical coupling and implications for other domain-swapped channels

When the first molecular structure of a eukaryotic voltage gated ion channel was determined, the S4-S5 linker from one subunit was found to contact the cytoplasmic half of S6 of the same subunit (11). This arrangement was made possible both by the bend in S6 at the conserved Pro-X-Pro motif, which curves S6_lower_ such that it is nearly parallel to the membrane surface, and the domain-swapped architecture of the channel. By studying chimeras of Shaker and KcsA, it was also known that the S4-S5 linker and S6_lower_ needed to be complementary for normal voltage dependent gating (32, 33). Based on these findings, the canonical model for voltage gated ion channels was proposed: upon hyperpolarization, the S4 moves intracellularly, which displaces the N-terminal end of the S4-S5 linker inwards, and that in turn compresses the S6 helical bundle by acting on S6_lower_. This model has found support in structures of mutant and cross-linked channels thought to mimic the hyperpolarized state, determined in membrane mimetics (23, 24, 34, 34–38).

Our findings portray a picture that differs substantially in the details. First, in the intermediate conformation, the S4 slides into an interfacial position past the S4-S5 linker without displacing it much. In this ‘down’ position it would apply a force on S5 of the adjacent subunit directly or through the S4-S5 linker (**Figure 3D, 5C**). Then in the constricted pore conformation, the S4 occupies a similar position but the N-terminal end of the S4-S5 linker is slightly displaced (by ∼2-3Å) inward and would also apply a force on S6_lower_ (**Figure 5B, C**). Thus, the voltage sensor movements are transmitted to the pore through two interfaces: one is the canonical interface involving same-subunit interactions and the second involves adjacent-subunit interactions where the S4 compresses the pore. Mutations at both interfaces have been shown to affect gating in different K_v_ channels (39, 40).

### A unifying mechanism for nondomain-swapped channels and domain-swapped channels

The above two-interface model describes how the voltage sensor regulates the pore in domain-swapped channels. What happens in nondomain-swapped channels? We have previously determined hyperpolarized and depolarized structures of the domain-swapped channel EAG1 and there found two interfaces for electromechanical coupling. In EAG1, the S4 in its down position moves past the S5 of the same subunit and compresses the pore while simultaneously pushing against the C-linker (cytoplasmic extension of S6) of the adjacent subunit (12). When one looks at the movements in K_v_2.1 (**Figure 6A**, also see **Movie S1** and **Movie S2**) and in EAG1 (**Figure 6B**), one realizes that domain-swapped and non-domain-swapped channels are quite similar. The principal difference between the two is simply whether each interface is formed with the same subunit or an adjacent subunit. Taken together, our results suggest a unifying mechanism for voltage-dependent gating in both domain-swapped and in non-domain-swapped channels. Perhaps this explains how a point-mutation in a TRP channel that changes its topology from domain-swapped to nondomain-swapped could result in a functional channel (41).

**Figure 6.**
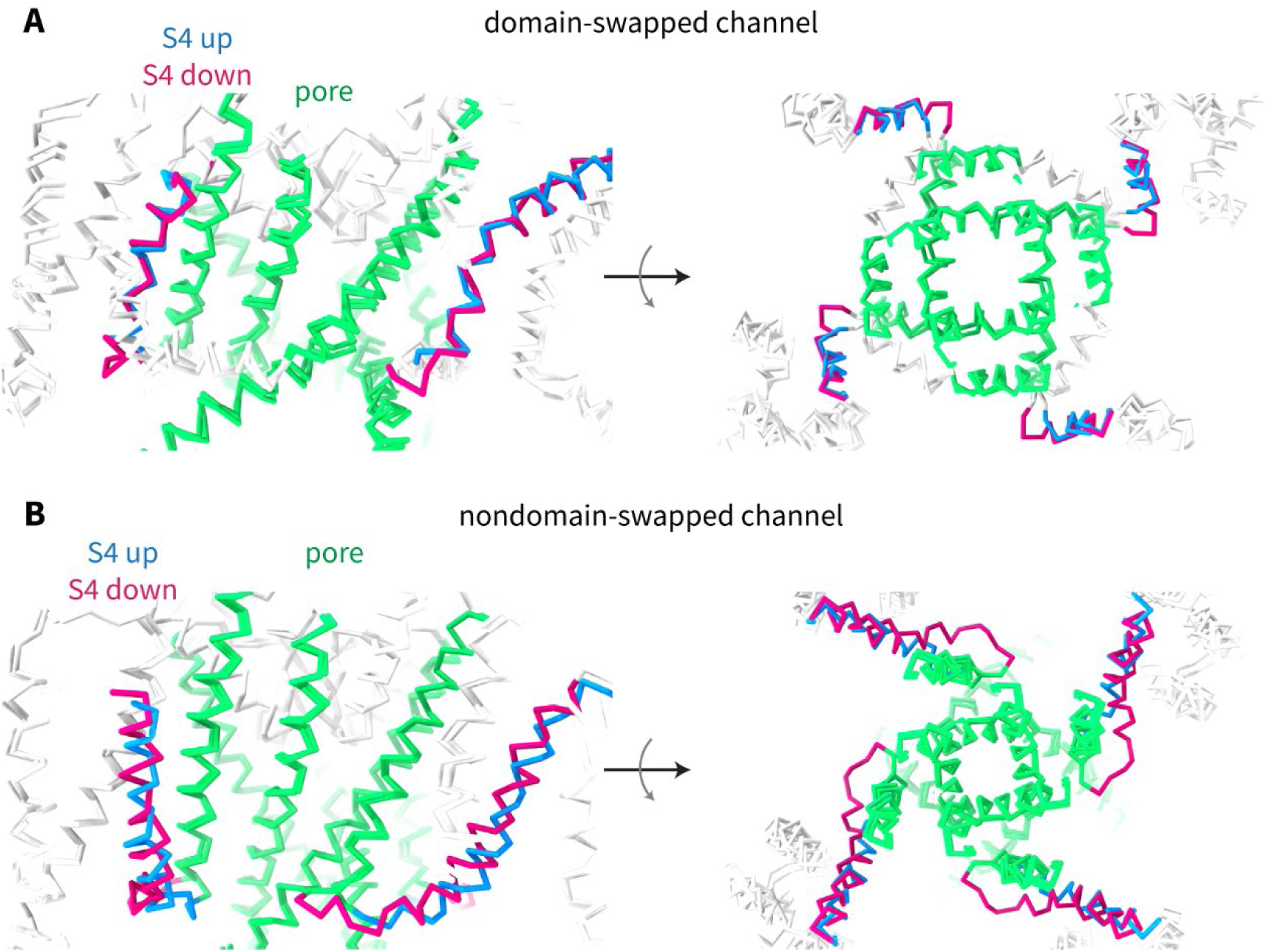
Voltage sensor movements in domain-swapped and nondomain-swapped channels. (**A-B**) Side views (*left*) and top-down views (*right*) of two conformations of the channel are shown for (**A**) domain-swapped K_v_2.1 and (**B**) nondomain-swapped EAG1 (PDB IDs: 8EOW and 8EP1). The pore of the channel is colored green, and the S4 helix in the voltage-sensor is colored blue for the up/depolarized conformation, and red for the down/hyperpolarized conformation.

### The mechanism of pore constriction: how does the gate in K_v_ channels work?

In our past experiments on hyperpolarizing EAG1 and KCNQ1, the gate was already locked shut by allosteric mechanisms – due to the presence of calmodulin in EAG1 (14) and the lack of PIP_2_ in KCNQ1 (42, 43). The structures of K_v_2.1 obtained here allow us to infer how the pore closes in a domain-swapped channel in which the voltage sensors exert force directly to gate the pore, with the limitation of medium resolution for the closed structure.

We find that the pore constricts by a rotation of the lower half of S6 with respect to the upper half of S6, centered around the second proline (P410) in the P-X-P hinge. The upper half of S6 stays in place, and it is unlikely that there are significant changes above the P-X-P motif. These observations are consistent with mutagenesis and electrophysiology efforts to characterize the gate of K_v_ channels, which found that: 1) the accessibility of residues both above and below the P-X-P motif do not change appreciably upon gate closure (44–46), and 2) mutation of either proline in the hinge to alanine results in a non-conducting channel (47, 48). Higher resolution structures are needed to detail the conformational changes further.

### Towards a description of the gating states in K_v_ channels

In our polarized cryo-EM dataset we use 3D classification to identify discrete states of K_v_2.1, but we do not know the electric field experienced by the proteins in each state. The fact that we observe different states, from the pore open conformation with the voltage sensors up, to the constricted pore conformation with the voltage sensors down, in the same dataset, indicates that we have a range of membrane potentials. While our experiment merely provides snapshots of different conformations sampled by K_v_2.1, electrophysiological studies of Kv2.1 have measured the voltage-dependence of S4 movement by recording gating currents (*blue* curve in **Figure 7A**) (20), and the voltage-dependence of pore opening and closing by recording ionic currents (*black* curve in **Figure 7A**) (49). We can thus compare the snapshots we observe in the cryo-EM experiment to the gating and ionic currents reported previously (20, 49) to infer what happens to a K_v_ channel, for instance, during the repolarization phase of an action potential (**Figure 7B**).

**Figure 7.**
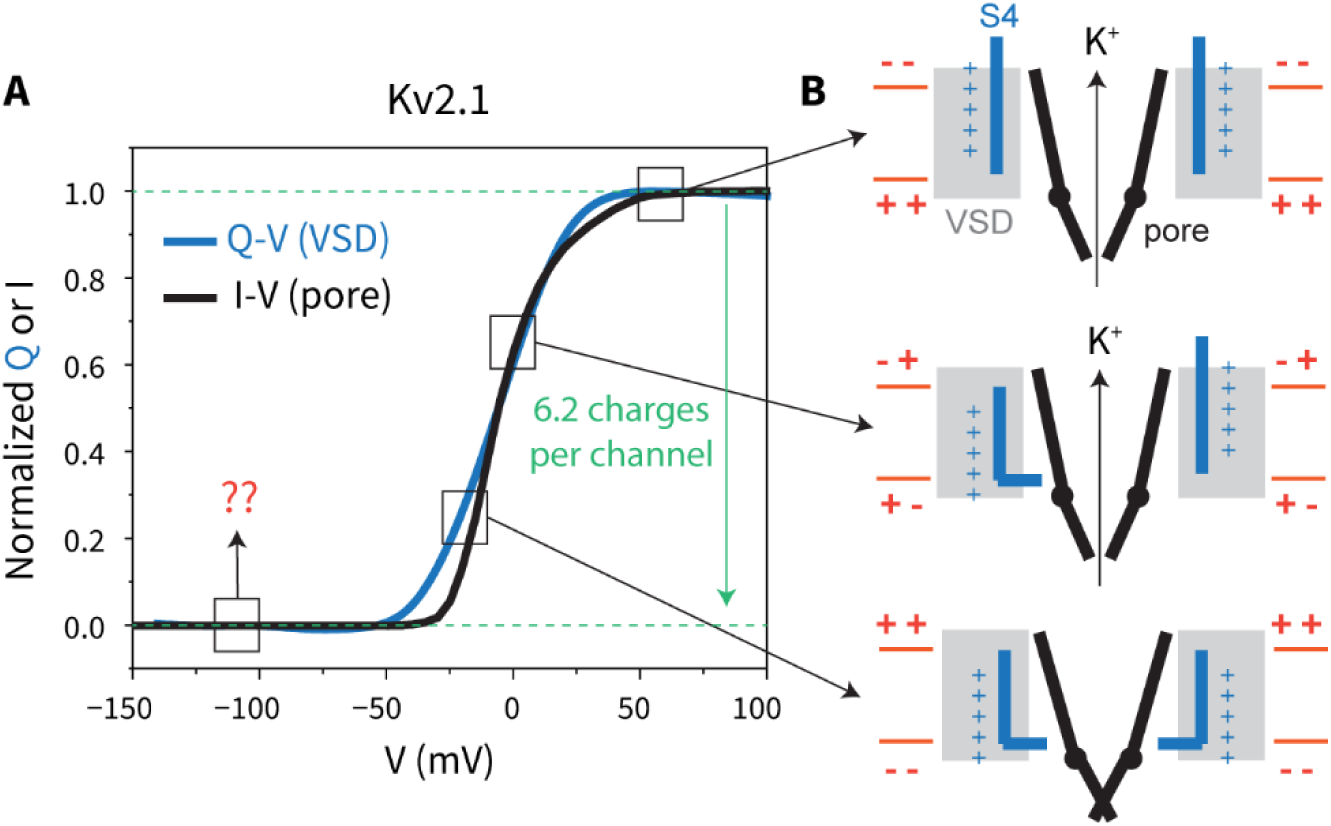
Gating cycle in a voltage-gated potassium channel. (**A**) Voltage-dependence of gating currents (Q-V, *blue*) and ionic currents (I-V, *black*) in K_v_2.1. The gating currents are from reference (20) and the ionic currents from reference (49). The total gating charge per channel was measured to be ∼6.2 elementary charges. (**B**) Cartoons showing the three conformations reported in this work: the depolarized conformation (*top*) with an open pore and all voltage sensors up, the intermediate conformation (*middle*) with an open pore but some voltage sensors down, and the constricted conformation (*bottom*) with a constricted pore and all four voltage sensors down. The S4 helix is colored blue, the VSD is colored light grey, and the pore is colored black, with only two subunits shown for clarity. It is possible that the voltage sensor in K_v_2.1 moves further than observed in this study; the location of such a state on the graph in panel A is marked by red question marks.

Starting from the depolarized state with all four voltage sensors up and the pore open (top cartoon in **Figure 7B**, the depolarized-lowK conformation), the flux of potassium ions from inside the cell to the outside begins to repolarize the membrane. The four voltage sensors in the channel begin to move as they sense the membrane potential, but up to ∼2 voltage sensors per channel can move to a 1-down position (one helical turn movement) while the most probable state of the pore is still open (middle cartoon, the intermediate conformation). Once ∼4 of the voltage sensors have moved to the 1-down (or further) position, then the pore is largely closed (bottom cartoon, the pore constricted conformation). The three conformations reported in this study are thus at least qualitatively consistent with measurements of gating charge and ionic currents in Kv2.1.

Finally, if the membrane potential becomes even more negative, the voltage sensors can keep moving to further down positions (the maximum gating charge per channel for K_v_2.1 is about 6.2 charges) (20, 21), with the pore remaining closed. We do not resolve structures of this hypothetical closed state for K_v_2.1 (red question marks in **Figure 7A**), but it would likely have all four voltage sensors moving down by ∼1-2 helical turns and a closed pore that might undergo further conformational changes. Elucidating these closed structures at even more hyperpolarized membrane potentials (∼-100 mV) would be particularly interesting in a channel such as Shaker, where the voltage sensors must move substantially further than those in K_v_2.1 (gating charge per channel of ∼13.5) (50–52), both to observe how S4 continues to move and to see what happens to the pore at even more negative voltages.

## Materials and Methods

### Cell lines

Sf9 (*Spodoptera frugiperda* Sf21) cells were used for production of baculovirus and were cultured in Sf-900 II SFM medium (GIBCO) supplemented with 100 U/mL penicillin and 100 U/mL streptomycin at 27 °C under atmospheric CO_2_.

HEK293S GnTl^−^ cells were used for protein expression and were cultured in Freestyle 293 medium (GIBCO) supplemented with 2% fetal bovine serum, 100 U/mL penicillin, and 100 U/mL streptomycin at 37 °C in 8% CO_2_.

### Expression and purification of K_v_2.1

Full-length human K_v_2.1 with a C-terminal GFP-His_6_ tag linked by a preScission protease (PPX) site was expressed as detailed before, using the BacMan system in HEK293S GnTI^-^ cells (13). The final SEC purification step used a Superose 6 Increase column (10/300 GL) pre-equilibrated with SEC buffer (10 mM Tris pH 8.0, 150 mM KCl (unpolarized) or 300 mM KCl (polarized), 0.03%:0.006% DDM:CHS, and 5 mM DTT). Fractions containing K_v_2.1 were pooled and concentrated at 2,000xg and 4°C to an A_280_ of ∼2 mg/mL.

### Reconstitution of K_v_2.1 into liposomes

Preparation of unpolarized liposomes containing K_v_2.1 was detailed before (13). In brief, purified K_v_2.1 channel was reconstituted into liposomes consisting of 90%:5%:5% POPC:POPG:cholesterol (wt/vol, Avanti Polar Lipids).(12, 13) Small unilamellar vesicles (SUVs) were produced by bath sonication in reconstitution buffer (10 mM Tris pH 8.0 and or 300 mM KCl (polarized)) and a low concentration (0.2% wt/wt for 10 mg/mL lipids) of the detergent C_12_E_10_ was added to the liposome suspension. Purified K_v_2.1 was mixed with lipids at a protein:lipid ratio of 1:20 (wt/wt) and detergent was removed over the course of ∼20 hours using 3 rounds of Bio-beads. The final concentration of lipids in the sample was 5 mg/mL.

Polarized vesicles were prepared as follows: 2 µM valinomycin (from an 8 mM stock in dimethyl sulfoxide) was added to the proteoliposomes and incubated for ∼30 minutes on ice. 70 µL of the vesicle solution was added to a 0.5 mL Zeba spin desalting column (40 kDa cutoff, Thermo Scientific), pre-equilibrated with sodium reconstitution buffer (10 mM Tris pH 8.0 and 300 mM NaCl), to exchange the external potassium for sodium. The sample was centrifuged at 1,500xg for ∼30 seconds at room temperature and ∼30 µL of flow-through containing vesicles was collected. The residual external K^+^ concentration is about 1 mM (12). 3.5 µL of the polarized vesicle solution was immediately applied onto a glow discharged Quantifoil R1.2/1.3 400 mesh holey carbon Au grid. After incubating the sample on the grid for 3 minutes at 20 °C with a humidity of 100%, the grid was manually blotted from the edge of the grid using a filter paper. Another 3.5 µL of the polarized vesicle solution was applied to the same grid for 20 seconds (53), and then the grid was blotted for 3 seconds with a blotting force of 0 and flash frozen in liquid ethane using a FEI Vitrobot Mark IV (FEI). Each grid with polarized vesicles used a freshly buffer exchanged sample.

During this procedure, the polarized vesicles are subject to a few minutes of incubation at room temperature followed by vitrification. It is likely that the vesicles that dissipate their ion gradients do so during this time.

### Cryo-EM data acquisition and processing

Data for the unpolarized K_v_2.1 liposomes were collected on a 300-keV FEI Titan Krios microscope located at the HHMI Janelia Research Campus. The microscope was equipped with a spherical aberration corrector (Cs corrector), a GIF BioQuantum energy filter and a Gatan K3 camera. A total of 17,007 movies were recorded on a single Quantifoil grid in super-resolution mode using SerialEM. The movies were recorded with a physical pixel size of 0.844 Å (super-resolution pixel size of 0.422 Å) and a target defocus range of - 1.0 to -2.0 µm. The total exposure time was ∼2 seconds (fractionated into 50 frames) with a cumulative dose of ∼60 e^−^/Å^2^.

Data for the polarized K_v_2.1 liposomes were collected on a 300-keV FEI Titan Krios 3 microscope located at the HHMI Janelia Research Campus. The microscope was equipped with a cold field emission gun (C-FEG), a Thermo Scientific Selectris X energy filter, and a Thermo Scientific Falcon 4i camera. A total of 20,339 movies were recorded on a single Quantifoil grid using SerialEM (54). The movies were recorded with a physical pixel size of 0.743 Å and a target defocus range of -0.8 to -1.8 µm. The cumulative dose was ∼60 e^−^/Å^2^.

The data processing workflow followed the same strategy previously reported for EAG1 and KCNQ1. Data processing was carried out using cryoSPARC v3/v4 (55) and RELION 4/5 (56). The movies were gain-normalized and corrected for full-frame and sample motion using the Patch motion correction tool (grid = 15×10). Contrast transfer function parameters were estimated from the motion-corrected micrographs using the Patch CTF estimation tool, which uses micrographs without dose-weighting. All subsequent processing was performed on motion-corrected micrographs with dose weighting. Particle picking was initially carried out using the Blob picker. 2D classes with clear protein density were used to train a TOPAZ picking model (57), which was then used to pick additional particles. Particles with clear protein density after 2D classification were pooled and duplicate picks were removed. An *ab initio* model was generated from 2D classes with clear secondary structure features and 3D classification and refinement was carried out either in cryoSPARC (Non-uniform refinement (58), Local refinement) or RELION (Blush regularization (59)). Symmetry expansion was carried out in RELION and symmetry-expanded particles were only subject to local angular searches during refinement.

### Model building and refinement

A structural model for the depolarized-highK conformation was built by docking four copies of the Alphafold-predicted (60) structure of K_v_2.1 into the up map (unpolarized dataset) and making adjustments as needed. The model was edited and refined using the ISOLDE (61) plugin in ChimeraX v1.5 (62) or WinCoot v0.98.1 (63) followed by real-space refinement in Phenix (64). The depolarized-lowK conformation, intermediate and pore constricted models were built starting from the depolarized-highK model, following a similar protocol. The quality of the final models was evaluated using the MolProbity (65) plugin in Phenix. Graphical representations of models and cryo-EM density maps were prepared using PyMOL (66) and ChimeraX (62). Map and structure statistics for the four structures are given in **Table S1** (**Supplementary Information**).

## Data Availability

The cryo-EM density map of K_v_2.1 from the unpolarized dataset has been deposited in the electron microscopy data bank under accession code EMD-XXXX and the corresponding model (depolarized-highK) has been deposited in the protein data bank under accession code XXXX. The depolarized, intermediate, and pore constricted maps from the polarized dataset have been deposited in the electron microscopy data bank under accession codes EMD-XXXX, EMD-XXXX and EMD-XXXX, respectively, and the corresponding models have been deposited in the protein data bank under accession codes XXXX, XXXX and XXXX, respectively.

## Author Contributions

V.S.M. performed the experiments. V.S.M. and R.M. designed the experiments, analyzed the data, and prepared the manuscript.

## Classification

Biological Sciences; Biochemistry.

## Acknowledgments

The authors would like to thank Mark Ebrahim, Johanna Sotiris, and Honkit Ng at the Evelyn Gruss Lipper Cryo-EM Resource Center for assistance with cryo-EM grid screening, Yi Chun Hsiung for assistance with insect and mammalian cell cultures, members of the MacKinnon Lab for helpful discussions, and Dr. Jue Chen and her research group for their input. We thank Rui Yan, Zhiheng Yu, and the rest of the staff at the HHMI Janelia CryoEM Facility for their support with cryo-EM microscope operation and data collection. Dr. Chia-Hseuh Lee carried out the cloning and initial biochemical characterization of the Kv2.1 channel. V.S.M. is supported by the Jane Coffin Childs Memorial Fund Fellowship. R.M. is an investigator in the Howard Hughes Medical Institute.

