## Supporting Information for "Electric field-induced pore constriction in the human Kv2.1 channel"

*Correspondence to: Roderick MacKinnon.

**This PDF file includes:**

Figures S1 to S5
Table S1
Legends for Movies S1 to S2

**Other supporting materials for this manuscript include the following:**

Movie S1
Movie S2

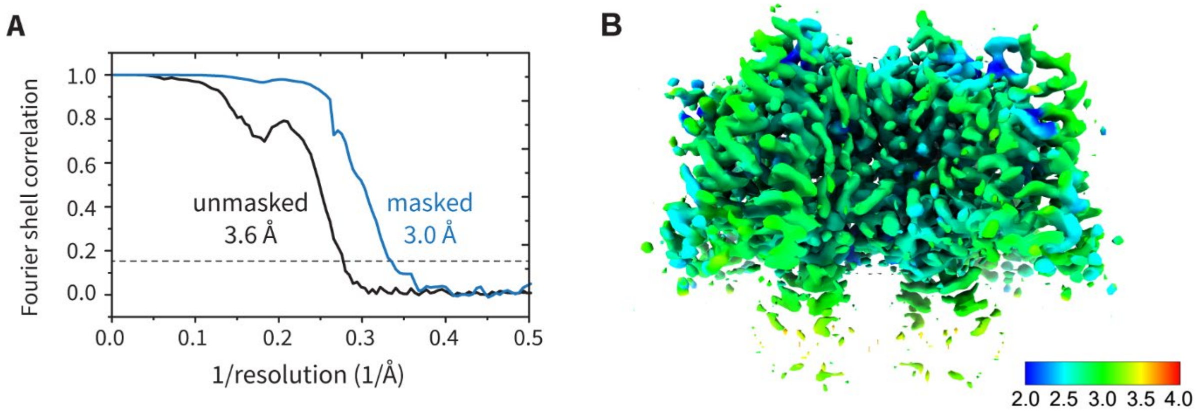


**Figure S1.** Fourier shell correlation curves and local resolution estimates for the depolarized map from the unpolarized dataset (i.e. the depolarized-highK conformation with all four voltage sensors up and an open pore).

(**A**) Fourier Shell Correlation (FSC) curves for the depolarized map from the unpolarized dataset calculated using the two independent half-maps from refinement. The FSC curve for the masked map is shown in *blue* and that for the unmasked map is in *black*. The nominal resolution at the gold-standard criterion (FSC=0.143, dashed *black* line) is indicated. (**B**) CryoSPARC-derived local resolution estimates (FSC = 0.143) for the same map overlaid with the sharpened map. The color key indicating the local resolution estimates is given in the bottom right corner.

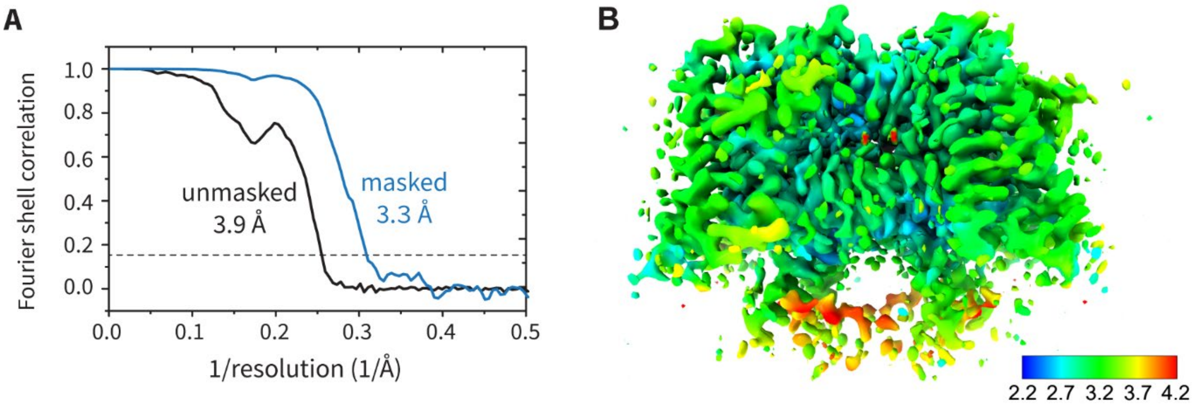


**Figure S2.** Fourier shell correlation curves and local resolution estimates for the depolarized map from the polarized dataset (i.e. the depolarized-lowK conformation with all four voltage sensors up and an open pore).

(**A**) Fourier Shell Correlation (FSC) curves for the depolarized map from the polarized dataset calculated using the two independent half-maps from refinement. The FSC curve for the masked map is shown in *blue* and that for the unmasked map is in *black*. The nominal resolution at the gold-standard criterion (FSC=0.143, dashed *black* line) is indicated. (**B**) CryoSPARC-derived local resolution estimates (FSC = 0.143) for the same map overlaid with the sharpened map. The color key indicating the local resolution estimates is given in the bottom right corner.

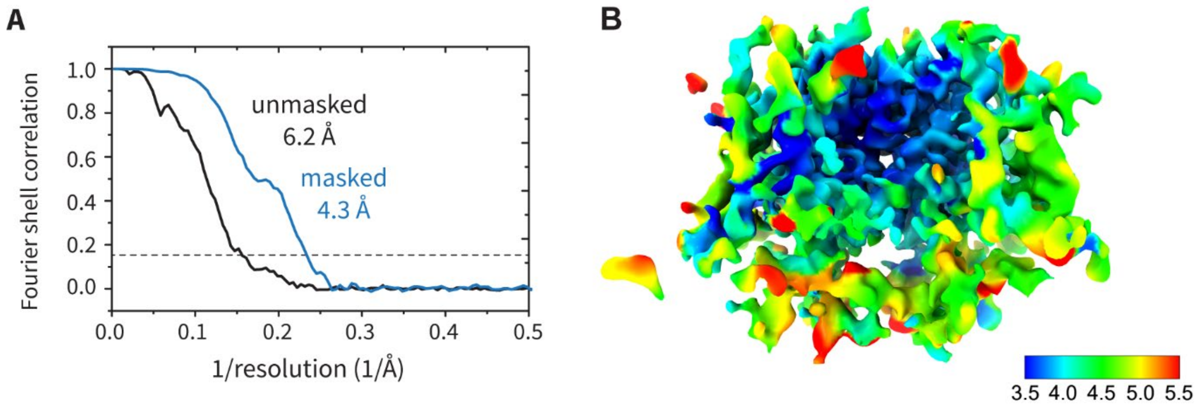


**Figure S3.** Fourier shell correlation curves and local resolution estimates for the hyperpolarized intermediate map from the polarized dataset (i.e. the conformation with 2 voltage sensors down and an open pore).

(**A**) Fourier Shell Correlation (FSC) curves for the hyperpolarized intermediate map from the polarized dataset calculated using the two independent half-maps from refinement. The FSC curve for the masked map is shown in *blue* and that for the unmasked map is in *black*. The nominal resolution at the gold-standard criterion (FSC=0.143, dashed *black* line) is indicated. (**B**) CryoSPARC-derived local resolution estimates (FSC = 0.143) for the same map overlaid with the sharpened map. The color key indicating the local resolution estimates is given in the bottom right corner.

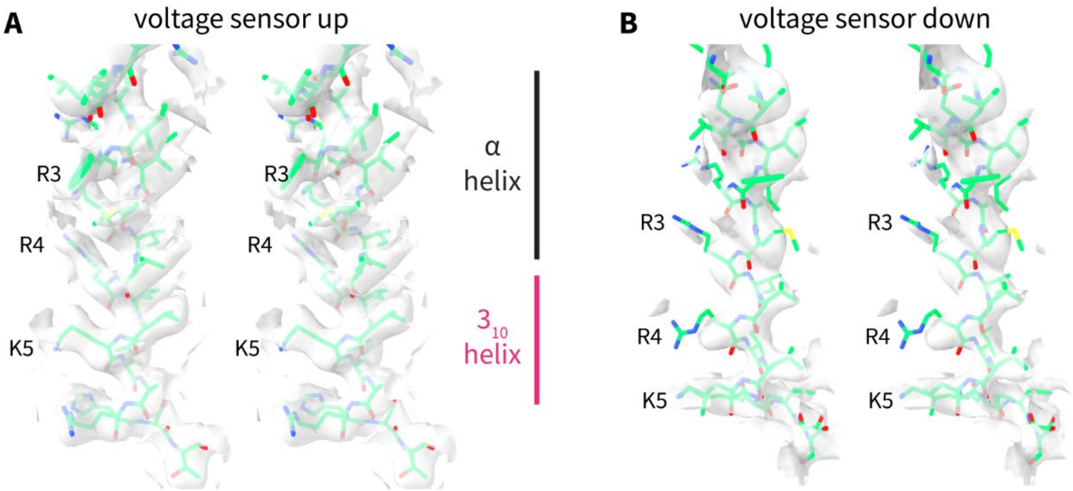


**Figure S4.** Comparison of the S4 helix in the voltage sensor in the up and down conformations.

(**A, B**) Stereoviews showing the S4 helix and associated cryo-EM density maps of the (**A**) depolarized conformation and the (**B**) hyperpolarized intermediate conformation, both from the polarized dataset. The helix is shown in *green* stick representation and the density map is shown as a translucent *grey* surface. The locations of three positive charges (R3, R4, and K5), and the α-helical and $3_{10}$-helical segments are indicated.

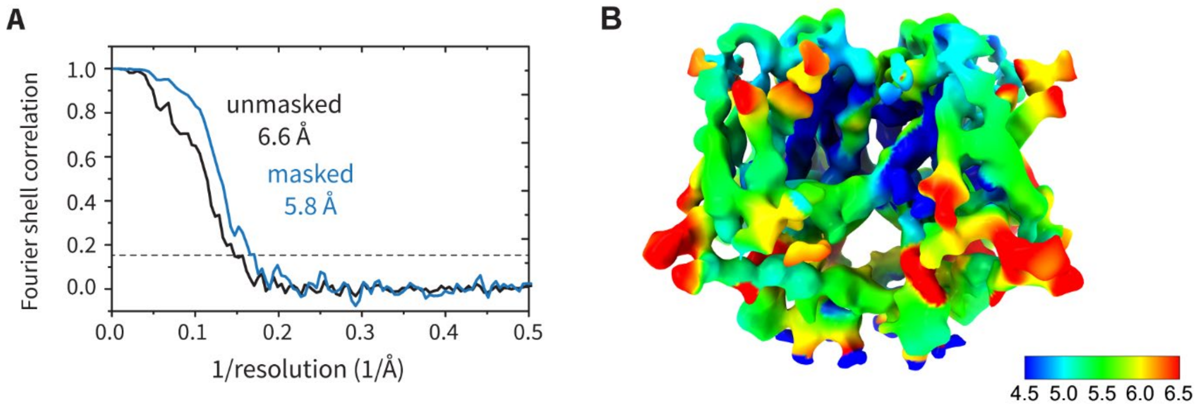


**Figure S5.** Fourier shell correlation curves and local resolution estimates for the hyperpolarized constricted map from the polarized dataset (i.e. the conformation with 4 voltage sensors down and a constricted pore).

(**A**) Fourier Shell Correlation (FSC) curves for the hyperpolarized constricted map from the polarized dataset calculated using the two independent half-maps from refinement. The FSC curve for the masked map is shown in *blue* and that for the unmasked map is in *black*. The nominal resolution at the gold-standard criterion (FSC=0.143, dashed *black* line) is indicated. (**B**) CryoSPARC-derived local resolution estimates (FSC = 0.143) for the same map overlaid with the sharpened map. The color key indicating the local resolution estimates is given in the bottom right corner.

**Table S1.** Summary of cryo-EM reconstruction and structural model statistics.

| **Reconstruction** | **Depolarized_highK** | **Depolarized_lowK** | **Intermediate** | **Constricted** |
|---|---|---|---|---|
| Dataset | Unpolarized | Polarized | | |
| Microscope/Camera | Titan Krios 2 300 kV, Gatan K3 | Titan Krios 3 300 kV, Falcon IVi | | |
| Pixel Size | 0.844 Å | 0.743 Å | | |
| Total dose | 60 e⁻/Å$^2$ | 60 e⁻/Å$^2$ | | |
| Defocus range | -1.0 to -2.0 µm | -0.8 to -1.8 µm | | |
| Movies collected | 17,007 | 20,339 | | |
| Particle number | 65,670 | 47,718 | 13,810 | 2,251 |
| Symmetry imposed | C4 | C4 | C1 | C4 |
| Nominal resolution (masked) | 3.0 Å | 3.3 Å | 4.3 Å | 5.8 Å |
| **Models** | **Depolarized_highK** | **Depolarized_lowK** | **Intermediate** | **Constricted** |
| **Ramachandran** | | | | |
| Preferred (%) | 98.47 | 97.97 | 95.66 | 96.91 |
| Allowed (%) | 1.53 | 2.03 | 4.34 | 3.09 |
| Outliers (%) | 0.00 | 0.00 | 0.00 | 0.00 |
| **MolProbity** | | | | |
| Clash Score | 4.31 | 3.15 | 4.20 | 4.90 |
| Rotamer Outliers (%) | 0.23 | 0.11 | 1.19 | 0.00 |
| Cβ deviations | 0.00 | 0.00 | 0.00 | 0.00 |
| Overall Score | 1.21 | 1.11 | 1.56 | 1.44 |
| **RMS deviations** | | | | |
| Bond lengths (Å) | 0.003 | 0.004 | 0.003 | 0.003 |
| Bond angles (°) | 0.46 | 0.49 | 0.47 | 0.50 |

**Movie S1 (separate file).** Sequence of conformational changes occurring during membrane hyperpolarization.

The movie shows a side view of the morph between two structures of $K_v2.1$: with the pore open and voltage sensors up (the depolarized-lowK conformation), and with the pore constricted and voltage sensors down (the hyperpolarized constricted conformation). The protein is shown in Cα trace representation. The S4 and S6 of one subunit are colored cyan and green, respectively. The Cα positions of positive-charged residues in the S4 helix are shown as blue spheres.

**Movie S2 (separate file).** Sequence of conformational changes occurring during membrane hyperpolarization.

The movie shows a top-down view (from the extracellular side) of the morph between two structures of $K_v2.1$: with the pore open and voltage sensors up (the depolarized-lowK conformation), and with the pore constricted and voltage sensors down (the hyperpolarized constricted conformation). The protein is shown in Cα trace representation. The S4 and S6 of one subunit are colored cyan and green, respectively. The Cα positions of positive-charged residues in the S4 helix are shown as blue spheres. The Cβ of P410 – the narrowest rigid constriction in the channel axis – is shown as orange spheres.
